# Tissue Folding by Mechanical Compaction of the Mesenchyme

**DOI:** 10.1101/164020

**Authors:** Alex J. Hughes, Hikaru Miyazaki, Maxwell C. Coyle, Jesse Zhang, Matthew T. Laurie, Daniel Chu, Zuzana Vavrušová, Richard A. Schneider, Ophir D. Klein, Zev J. Gartner

## Abstract

Many tissues fold during development into complex shapes. Engineering this process in vitro would represent an important advance for tissue engineering. We use embryonic tissue explants, finite element modeling, and 3D cell patterning techniques to show that a mechanical compaction of the ECM during mesenchymal condensation can drive tissue folding along programmed trajectories. The process requires cell contractility, generates strains at nearby tissue interfaces, and causes specific patterns of collagen alignment around and between condensates. Aligned collagen fibers support elevated tensions that promote the folding of interfaces along paths that can be predicted by finite element modeling. We demonstrate the robustness and versatility of this strategy for sculpting tissue interfaces by directing the morphogenesis of a variety of folded tissue forms from engineered patterns of mesenchymal condensates. These studies provide insight into the active mechanical properties of the embryonic mesenchyme and establish entirely new strategies for more robustly directing tissue morphogenesis *ex vivo,* without genetic engineering.

## INTRODUCTION

Engineered tissues have applications in basic sciences, drug testing, and regenerative medicine (Bajaj et al., 2014; Clevers, 2016; Huch et al., 2017). A key challenge for tissue engineers is to build or grow tissues in vitro that reproducibly incorporate key structural motifs of the corresponding tissue in vivo (Gjorevski et al., 2016; Huh et al., 2010; Lancaster and Knoblich, 2014; Lancaster et al., 2017; Warmflash et al., 2014). Tissue folds are a widespread and crucially important structural motif because they contribute to tissue function in adults. However, the trajectory of folding during development also contributes to changes in the detailed structure of a tissue, through processes such as cell identity specification and the emergence of anisotropies in the distribution of cells and extracellular matrix fibers (Kim et al., 2015; Li et al., 2017; Shyer et al., 2015). Thus, the gross folded form of a tissue, as well as the trajectory of tissue folding, can both be important for mature tissue function.

While tissue folding in vivo can be highly reproducible and robust (Nelson, 2016; Savin et al., 2011), folding remains difficult to reconstitute or control in vitro (Huch et al., 2017; Li et al., 2017; Varner and Nelson, 2014b). For example, many popular organoid models can faithfully reproduce epithelial architecture and composition across tens to hundreds of microns, but the size and shape of folded architectures can vary considerably from organoid to organoid, particularly over larger length-scales (Huch et al., 2017; Takebe et al., 2015). Folding is difficult to control in vitro because it emerges from spatially patterned cell dynamics that span micro- to millimeter scales. Moreover, folding is critically reliant on boundary conditions and often requires interactions between more than one tissue layer (Nelson, 2016; Sydney Gladman et al., 2016). In principle, engineers could guide the formation of folds from simpler structures by combining top-down engineering with the intrinsic self-organizing properties of cells and extracellular matrix. According to this strategy, engineers would use top-down tools to define the initial conditions of a culture (Qi et al., 2013; Wang et al., 2017; Zhang and Khademhosseini, 2017), and then leverage the autonomous cellular processes underlying tissue self-organization to drive folding along a single trajectory.

To implement such a strategy for engineering the formation of precisely folded tissues, we considered the physical mechanisms through which folding occurs in living systems. Folds form when initially planar regions of tissue bend or buckle to form more complex 3D shapes. Here, the concave surface of a tissue layer experiences negative strain (reduction in dimension) and the convex surface experiences positive strain (increase in dimension), Figure 1A (Timoshenko and Woinowsky-Krieger, 1959). Tissue folding therefore derives from a mismatch in strains between two adjacent tissue layers, which in vertebrate embryos are often an epithelium and the underlying extracellular matrix (ECM) in a loose connective tissue layer known as the mesenchyme (Hay, 2013). The requisite strains arise in response to stresses (force per unit cross-sectional area) that accumulate during developmental processes. For example, compressive or tensile stresses can act across entire organ systems to generate patterns of folding through buckling processes (Nelson, 2016; Shyer et al., 2013), or they can act across smaller regions of a tissue, for example when smaller groups of cells proliferate or exert cytoskeletal forces locally against surrounding tissue (Kim et al., 2015; Mammoto et al., 2013; Odell et al., 1981; Panousopoulou and Green, 2016; Wen et al., 2017). While both classes of folding are amenable to engineering, we focused on mechanisms that act at the level of local cell-generated forces. These mechanisms are readily integrated with top-down patterning technologies such as optogenetics, micro-molding, and printing approaches that control cellular and ECM tissue composition at specific locations (Bhattacharjee et al., 2015; Hinton et al., 2015; Miller et al., 2012; Oakes et al., 2017; Stevens et al., 2013; Warmflash et al., 2014).

**Figure 1.**
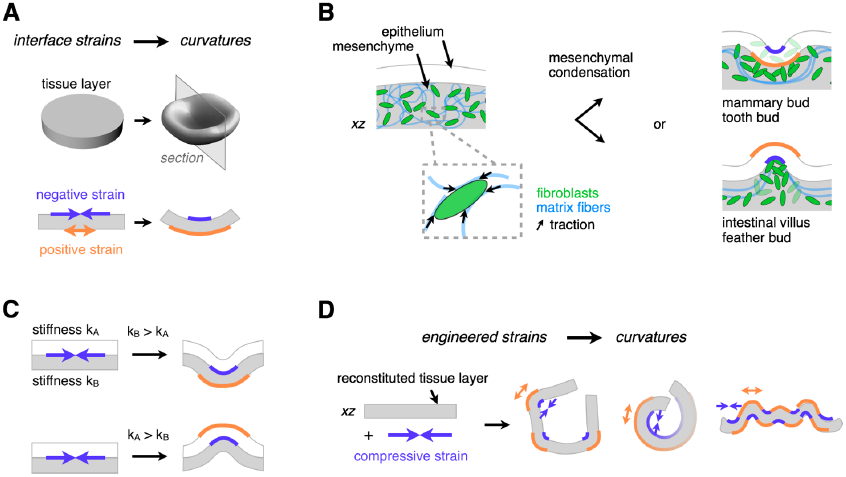
Tissue Folding Requires Patterns of Negative and Positive Strain at Tissue Interfaces. (A)Curvature at tissue interfaces is associated with a decrease in length (negative strain) at concave interfaces, and an increase in length (positive strain) at convex interfaces. (B)Strains at the epithelial-mesenchymal interface are associated with folding events in the mammary bud, tooth bud, intestinal villi, and feather bud. Strained regions of tissue occur adjacent to mesenchymal condensates in these tissues. As mesenchymal cells generate traction forces on nearby ECM, they could participate in driving such folding events. (C)Negative strain at the interface between two tissue layers can drive invagination or evagination according to the relative bending stiffnesses of the adhered layers.(D)Patterns of mesenchymal condensates could be mapped to corresponding patterns of tissue strains, creating a programmable approach to engineered folding of reconstituted tissues.

In order to identify cellular processes that could be used to direct tissue folding downstream of patterning technologies, we looked to developmental systems where the pattern of folding is tightly linked to patterns of cell dynamics. For example, the pattern of folds that emerge during the morphogenesis of teeth (Mammoto et al., 2011), mammary buds (Cowin and Wysolmerski, 2010; Sakakura et al., 1976), feathers (Eames and Schneider, 2005; Jiang et al., 1999; Wessells and Evans, 1968), gut villi (Walton et al., 2012), salivary glands (Hieda and Nakanishi, 1997), and lung buds (Kim et al., 2015) are all spatially and temporally coupled to corresponding patterns of cellular condensates in the neighboring mesenchyme (Figure 1B). In particular, experiments using explants of the developing mouse intestine suggest that mesenchymal condensates forming at the epithelial-mesenchymal interface contribute to the early stages of the folding of the gut to form villi, because they can be physically isolated from surrounding tissue without disrupting the initiation of local folds (Walton et al., 2016b). Similar observations have also been made for feather bud formation during culture of chick skin (Davidson, 1983). We hypothesized that mechanical strains sufficient to produce local curvature at these interfaces could be generated directly by mesenchymal condensation, since local tissue mechanical properties change at the site of condensation (Mammoto et al., 2011). In this model, invagination (inwards bending) or evagination (outwards bending) would arise from the relative stiffness of the two conjoined layers (Oster and Alberch, 1982). More specifically, folding would occur towards the tissue layer with the lower bending modulus (Figure 1C, Figure S1).

Here, we develop an engineering framework for guiding autonomous tissue folding along prescribed trajectories by using patterns of mesenchymal condensates to program corresponding patterns of strains into slabs of ECM gels (Figure 1D). We begin by providing evidence that cell dynamics associated with mesenchymal condensation can bend nearby epithelial-mesenchymal interfaces. Specifically, we observe that mesenchymal condensation in the developing mouse intestine and chick skin is associated with non-muscle myosin II-dependent compaction of the collagenous ECM. This compaction is associated with alignment of nearby collagen fibers, and tissue interface curvature. Moreover, we demonstrate through in vitro reconstitution that mesenchymal condensation is sufficient to generate these in vivo hallmarks in reconstituted cell/ECM gel composites. To place tissue folding by mesenchymal condensation under engineering control in vitro, we perform quantitative measurements relating initial cell patterns to tissue strains and interface curvatures, allowing folding trajectories to be predicted through finite element modeling. Finally, we utilize this system to drive the folding of complex 3D structures that mimic in vivo-relevant and other, more stylized folding motifs.

## RESULTS

### Signatures of Mesenchymal Condensation Mechanics

We first sought to identify patterns of cell dynamics that could be leveraged to drive folding of reconstituted tissues. We first focused on embryonic villus formation in the mouse gut. Previous reports suggest that condensates in the embryonic intestine are tightly associated with the folding of the epithelial-mesenchymal interface (Walton et al., 2012; 2016b). Consistent with these reports, we found that folding initiated in the mouse gut at the basal surface of the epithelium, and was tightly associated with the appearance of clusters of PDGFR+ mesenchymal fibroblasts. Collagen I accumulated around these PDGFR+ foci during folding. Importantly, cell condensation, collagen accumulation, and tissue folding were blocked by treatment with blebbistatin, an inhibitor of myosin II activity (Fig. 2A-C, Figure S2A-C). Similar results were observed for feather bud formation in chick skin (Figure S2D-H). These observations are consistent with a role for mechanical condensation of the mesenchyme along the epithelial-mesenchymal interface in driving the folding of nearby tissue interfaces. They further suggest that if reconstituted in vitro, the mechanics of the condensation process could be leveraged to direct the local folding of tissue interfaces.

**Figure 2.**
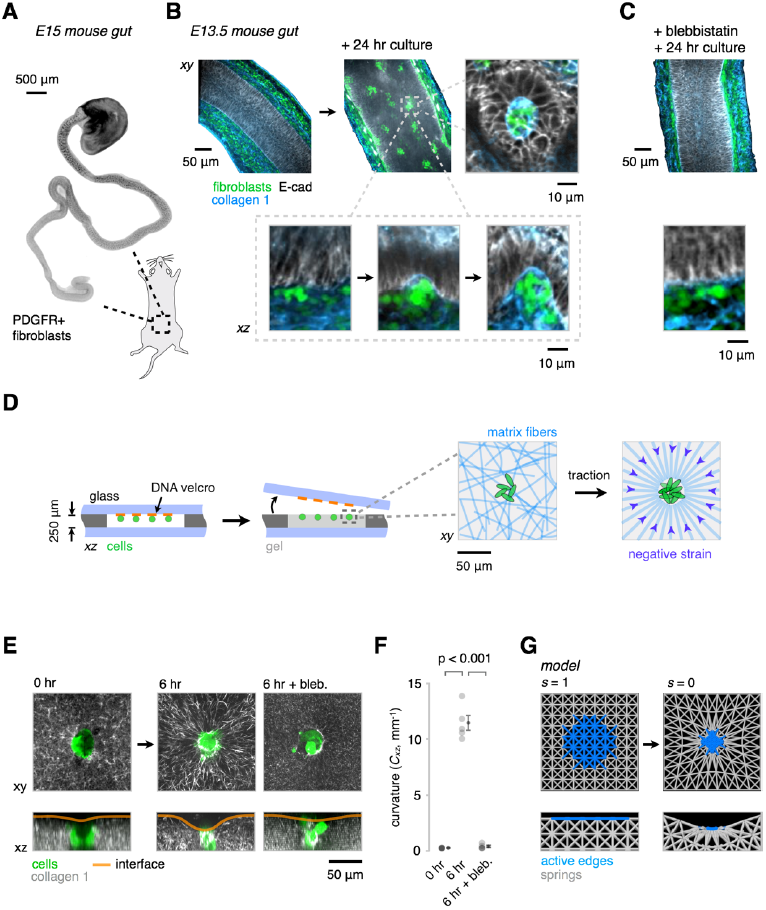
Signatures of Mesenchymal Condensation in the Mouse Gut, and their Reconstitution In Vitro and In Silico. (A)Whole-mount wide-field fluorescence microscopy image of the embryonic day 15 (E15) mouse intestine showing PDGFR+ fibroblast clusters forming in an anterior to posterior wave. (B)Optical sections from whole-mount confocal immunofluorescence images showing PDGFR+ cells (green) and collagen I fibers (blue) in E13.5 explants cultured for 0 or 24 hr in vitro. Detail shows intermediate stages of PDGFR+ cluster formation against the basal surface of the epithelium (E-cadherin, gray), along the wave of condensation. Successive stages of cluster formation show progressive collagen I accumulation and localized curvature at the basal surface of the epithelium. (C)Intestine explant cultures as in (B) show reduced cell clustering, collagen I accumulation, and interface curvature in the presence of 30 μ M blebbistatin (a myosin II inhibitor). See Figure S2A-C for further quantitation. (D)Schematic of reconstitution strategy using DNA-programmed assembly of cells (DPAC) to build loose clusters of mesenchymal cells near the surface of ECM gels containing collagen I and matrigel. Detail at right illustrates the hypothesized traction-mediated compaction and alignment of ECM fibers around cell clusters. (E)GFP-expressing MEF clusters (green) were patterned in AF555-labeled collagen I-containing gels (gray) as in (D). Live confocal microscopy of condensing clusters and collagen I reveals ECM compaction, radial collagen I fiber alignment, and the emergence of curvature of the gel-medium interface. These phenomena are blocked by treatment with 30 μ M blebbistatin. (F)Quantification of the interfacial curvatures proximal to the condensates shown in (E) (mean ± SEM, *n* = 5, one-way ANOVA with Holm-Sidak’s multiple comparisons test). (G)Snapshots from a finite-element model containing passive elastic elements (gray) and active edges (blue) whose length *s* can be reduced to simulate local gel strains by cell clusters.

### In Vitro Reconstitution of Mesenchymal Condensation Mechanics

To more rigorously investigate whether the cell dynamics observed in vivo were sufficient to direct tissue folding, as would be necessary to place the folding process under engineering control, we reconstituted a simulacrum of the mesenchymal-epithelial tissue interface, devoid of the overlying epithelium. We used a 3D cell patterning technology called DNA-Programmed Assembly of Cells (DPAC) to prepare loose clusters of GFP-expressing mouse embryonic fibroblasts (MEFs) positioned just below the surface of 250 µm-thick slabs of fluorescently-labeled collagen I in matrigel (Figure 2D) (Todhunter et al., 2015). We attached these slabs to glass coverslips and tracked cell and ECM dynamics within them using time-lapse confocal microscopy. Previous studies have shown that cells in these types of gels generate traction forces that strain and align matrix fibers (Baker et al., 2015; Harris et al., 1981; Sawhney and Howard, 2002; Vader et al., 2009). Consistent with these reports, MEF clusters compacted over six hours in culture, concentrating collagen near the cluster surface, and producing patterns of radially aligned collagen fibers in their local microenvironment (Figure 2E, Figure S3A-E, movie S1). Examination of the gel-media interface revealed local curvature and invagination of the ECM gel proximal to the condensing MEFs. This curvature motif was similar, but of opposite polarity to that seen in the mouse gut, possibly due to the absence of a stiff overlying epithelium (Figure S1). As in the intestinal gut explants, cluster compaction, collagen alignment, and interfacial curvature were significantly inhibited by treatment with blebbistatin (Figure 2E,F). These observations suggest that key features of condensation can be reconstituted in vitro, and that condensates themselves could act as mechanical actuators of tissue interface curvature (Oster et al., 1983). Indeed, a two-parameter finite element model (FEM) consisting of an isotropic contractile node within a grid of unit cells constructed from elastic springs captured aspects of elastic edge (ECM fiber) compaction, alignment, and interface invagination seen in the in vivo and in vitro systems (Figure 2G).

ECM compaction and cell motility have both been implicated in mesenchymal cell condensation (Mammoto et al., 2011; Oster et al., 1983). While both processes could contribute to condensation, ECM compaction among them could account for strains at nearby tissue interfaces sufficient to explain their folding. To evaluate the relative contribution of cell motility and ECM compaction to the condensation process in vitro, we prepared circular grids of MEFs at the ECM surface using DPAC (Figure 3A). We then measured the radial strain (shrinkage) of the cell grid by tracking individual cells, and of the ECM by tracking fluorescent collagen I fibers dispersed in the underlying matrix (Figure 3B). We found that the cell grid condensed towards a central focus, and that the total radial strain was primarily accounted for by strain in the ECM, rather than by migration of cells through the ECM towards the focus (Figure 3C). Further, the condensation process was abolished by inhibiting myosin II with blebbistatin. These data are consistent with a model wherein reconstituted cell condensates act as mechanical actuators by compacting the surrounding ECM and imposing local strains at tissue interfaces.

**Figure 3.**
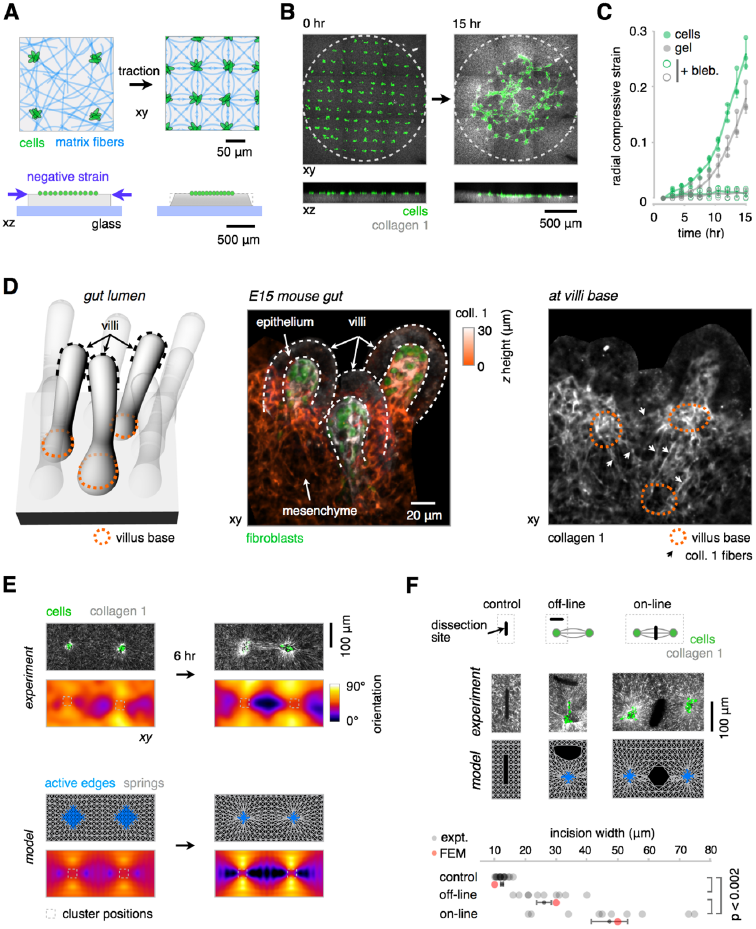
Mechanical Coupling Between Mesenchymal Condensates In Vivo and In Vitro. (A)Schematic for measuring cell migration and ECM compaction contributions to reconstituted tissue condensation. A grid of GFP-expressing MEF clusters (green) is patterned just below the surface of a gel containing collagen I (gray), and the motion of fluorescent collagen fibers and cells are tracked by time-lapse confocal microscopy. (B) Maximum intensity projections and xz sections from a representative time-lapse confocal microscopy experiment showing MEF cluster grid (green) and AF555-labeled collagen I (gray) converging towards a central focus. The dotted lines mark the initial spatial extent of the grid. (C)Quantification of the data in (B), showing radial strain of the cell grid (green) and ECM (gray) in the presence and absence of 30 μ M blebbistatin (bleb., mean ± SEM, *n* = 3). (D)(Left) Schematic illustrating the developing mouse intestine shown in the images to the right. Dark dotted lines indicate the location of three villi caps, and orange dotted lines indicate the position of three villi bases. (Center) Immunofluorescence image of the embryonic day 15 (E15) mouse intestine showing three representative villi as in the diagram to the left. Data is shown as a maximum intensity projection of stained collagen I fibers within these villi, color coded according to their height in the confocal stack. PDGFR+ fibroblast clusters (green) are also shown. (Right) One section from the base of the stack illustrating regions of collagen I alignment (white arrows) around villi bases (orange dotted lines). (E)(Top) GFP-expressing MEF clusters (green) were patterned using DPAC at fixed distances within a gel containing collagen I (gray). Collagen alignment is evident after 6 hr; the pixel-wise angle of average orientation was used to generate the heat-maps below each microscopy image. (Bottom) Qualitatively similar patterns of elastic edge alignment (gray) were observed between contractile nodes (blue) in the finite-element model (FEM). (F)Retraction of the gel surface after laser ablation in three regions around pairs of interacting condensates. Qualitatively similar retraction behavior in the FEM implies that collagen straps bear greater tension than the less aligned regions of the gel (mean ± SEM, *n* > 9 incisions per group, one-way ANOVA with Holm-Sidak’s multiple comparisons test).

### Strain Engineering among Mechanically Coupled Arrays of Condensates

Condensates occur in arrays in a variety of embryonic systems, where they can become mechanically coupled through networks of aligned ECM fibers. For example, we observed distinct patterns of collagen alignment between adjacent feather buds in chick skin (confirming previous reports (Stuart and Moscona, 1967), Figure S2H), and additionally observed aligned collagen fibers spanning the basal plane of nascent villi in the developing mouse intestine (Figure 3D). We hypothesized that these regions of aligned matrix fibers were a consequence of the condensation process, and were under elevated tension. Moreover, we reasoned that if similar patterns could be reconstituted in vitro, they could be used to coordinate tissue folding over much larger distances and into complex three-dimensional geometries.

To explore the mechanical coupling between networks of condensates, we used DPAC to prepare pairs of loose MEF clusters at set distances from one another. As the paired clusters condensed and remodeled the surrounding ECM over 6 hr, collagen fibers were initially radially aligned around each condensate, and then formed regions of amplified collagen fiber alignment (“straps”) along the axis between them (Figure 3E, Figure S3F-J). These straps were similar to the aligned collagen tracts seen in chick and mouse tissues (Figure 3D, Figure S2, movie S2) (Sawhney and Howard, 2002). A FEM simulation of this process qualitatively captured the same pattern of aligned elastic elements between nodes and suggested that these regions coincided with higher tensile stresses relative to regions having less strongly-aligned collagen fibers. To evaluate whether straps were under elevated tension compared to other regions of the gel, we mapped the local distribution of forces around interacting condensates by measuring recoil of the ECM surface upon cutting. Biophysical studies suggest that regions of gel under elevated tensile stresses should undergo a damped elastic recoil in proportion to the local tension orthogonal to the incision (Bonnet et al., 2012; Kumar et al., 2006; Legoff et al., 2013), (Method Details). Indeed, in both experiments and FEM, significantly greater recoil of the gel surface occurred upon ablation with a focused UV laser at straps in comparison to regions having only radially aligned collagen or to control regions distant from condensate pairs (Figure 3F). These data suggest that nearby condensates are mechanically coupled, and can promote the alignment of collagen fibers along their axes of interaction. Moreover, the data suggest that these aligned collagen fibers are under elevated tension compared to the surrounding tissue.

From an engineering perspective, the same phenomena that drive changes in curvature around individual cell clusters could be harnessed to program the curvature of deformable tissue interfaces across considerably larger distances. In such a scenario, the positions and density of condensing mesenchymal cells would determine tension patterns borne by aligned collagen straps. Without physical constraints on the geometry of the gel, these tensions would drive patterns of interfacial strains, thereby having a direct, predictable, and reproducible relationship to the final folded architecture of the tissue. However, implementing such a strategy would require that straps among patterned condensates formed in a predictable pattern, for example between nearest neighbors, similar to the patterns observed among feather buds in the developing chick.

We explored how networks of condensates mechanically couple by studying the relationship between the initial pattern of interacting condensates, and the resulting pattern of force-transmitting collagen straps. We again used DPAC to place loose clusters of MEFs in either isotropic grids (with the same densities along the horizontal and vertical axes, ρ_x_ and ρ_y_) or anisotropic grids (unequal ρ_x_ and ρ_y_) at the surface of ECM gels adhered to the culture surface. We found that collagen straps formed preferentially between nearest neighbor clusters as they condense (Figure 4A). Straps persisted in orientation for as long as neighboring aggregates did not merge with each other during the condensation process, suggesting that the spatial pattern of forces imposed at reconstituted tissue interfaces could be predictably encoded by the geometry of initial MEF cluster positions.

**Figure 4.**
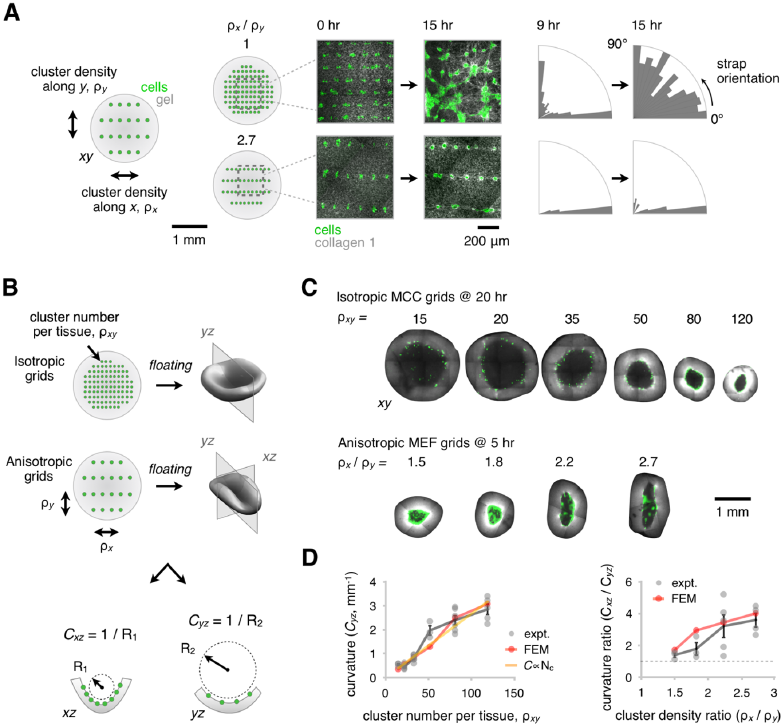
Patterns of Condensates Mechanically Couple over Large Distances and Quantitatively Encode the Trajectory of Tissue Curvature. (A)(Left) Isotropic and anisotropic grids of GFP-expressing MEF clusters (green) were patterned using DPAC into gels containing AF555-labeled collagen I (gray), adhered to a glass substrate, and imaged using time-lapse confocal microscopy. The density of clusters along the horizontal and vertical axes are denoted p_x_ and p_y_, respectively. The emergence of collagen straps (center) was observed over 9 to 15 hr. (Right) Radial bar charts illustrating the alignment of tension-bearing straps relative to the horizontal axis at 9 and 15 hr. Straps extend only between nearest neighbors in the vertical and horizontal directions, until later timepoints when the clusters begin to merge. (B)Schematic depicting how curvature is measured in the xz and yz planes of floating gels containing either isotropic cell cluster grids (with total cluster density p_xy_) or anisotropic grids. (C)Representative confocal microscopy sections through the mid-plane of invaginating reconstituted tissues patterned with different total cluster densities and combinations of densities in the horizontal and vertical directions. Graded patterns of curvature emerge for the indicated density and anisotropy of cell clusters. (D)Curvature measurements for the experimental data shown in (C). (Left) Curvature as a function of total cluster density and (right) curvature anisotropy as a function of anisotropy in cluster density along *x* and *y*. These data constitute quantitative calibration relationships (mean ± SEM, *n* > 2 per grid geometry) that adequately parameterized a finite element model (FEM) relating “blueprint” patterns of contractile nodes to folding trajectories of reconstituted tissues.

### Engineering Control of Interfacial Curvature in Reconstituted Tissues

Given that strains at tissue interfaces can be relieved by out-of-plane buckling (Armon et al., 2011), we hypothesized that grids of condensing MEFs would drive reconstituted tissue folding if they were released from the geometric constraints imposed by the glass substrate. We therefore studied the global curvatures of MEF grids placed at the surface of free-floating, rather than substrate-anchored ECM gels (Figure 4B). We found that gels followed predictable trajectories as they underwent morphogenesis from planar to curved 3D geometries. Specifically, gel layers carrying grids of loose cell clusters formed radially-symmetric invaginations (Figure 4C). Moreover, by patterning a range of cell types with different contractile properties, we found that the initial curvature rates of the invaginating gels were proportional to the rates at which the cells strained the surrounding ECM, with curvature dynamics spanning timescales of ~ 5 to > 40 hours (Figure S4A-E).

We chose relatively slower-folding reconstituted tissues bearing isotropic grids of mesenchymal-like mammary carcinoma cells (MCCs) to enable detailed measurement of the temporal dynamics of folding by confocal microscopy. We found that the gel curvature *C* at a given time-point increased approximately linearly with the cell cluster number per tissue (ρ_xy_, Figure 4C,D). The experimental *C* vs. ρ_xy_ data were adequately fit by both a proportional scaling relationship describing bending induced by constant strain-rate actuators (Holmes et al., 2011; Timoshenko and Woinowsky-Krieger, 1959) (R^2^ = 0.8, Method Details), and by a 3D implementation of our FEM. We additionally found that the emergence of curvature coincided with extensive and irreversible gel remodeling since pre-folded tissues unfolded by less than 40% upon inducing apoptosis of cells using staurosporine (Figure S4F,G).

Beyond radially symmetric invaginations, we predicted that anisotropic grids of MEF clusters having a greater density along the horizontal relative to the vertical axis (ρ_x_ > ρ_y_) would program a fold running vertically, since tension-bearing collagen straps would be concentrated along the more densely packed horizontal axis. Indeed, anisotropic MEF grids consistently folded gels along the expected axis, and curvature anisotropy in *x* and *y* was approximately proportional to cluster density anisotropy (Figure 4C,D). Surprisingly, identical grids actuated by less contractile and more migratory MCCs did not form controlled anisotropic folds along a predictable axis (Figure S4H). We found that the ratio of folding rate to cell migration rate for a given condensing cell type was critical in determining the fidelity with which anisotropic folds could be specified, because rapidly migrating cells quickly scramble the initial cluster pattern imposed by DPAC (see Method Details, Figure S4I,J).

We continued to explore the generality and robustness with which different patterns of mesenchymal condensates could encode the morphogenesis of reconstituted tissues into a variety of complex 3D folding motifs (Figure 5, movie S3, Figure S5). These included the isotropic and anisotropic motifs already discussed, as well as a compound curvature motif (spatially graded curvature) and an opposing curvature motif (neighboring tissue regions with curvatures of opposite polarity, Figure S5A). We first elaborated on the anisotropic folding motif to generate a coiled tissue conceptually similar to several biological forms, including the looping of the small intestine. To drive coiling, we placed anisotropic grids of MEF clusters at a 45° angle to the long axis of a rectangular ECM substrate, encoding a partially-enclosed helical shape with a pitch of 1 mm and radius of 200 microns (with comparable values of 1.2 mm and 220 microns in a member of the corresponding FEM family, Figure 5A,B, Figure S5B,C).

**Figure 5.**
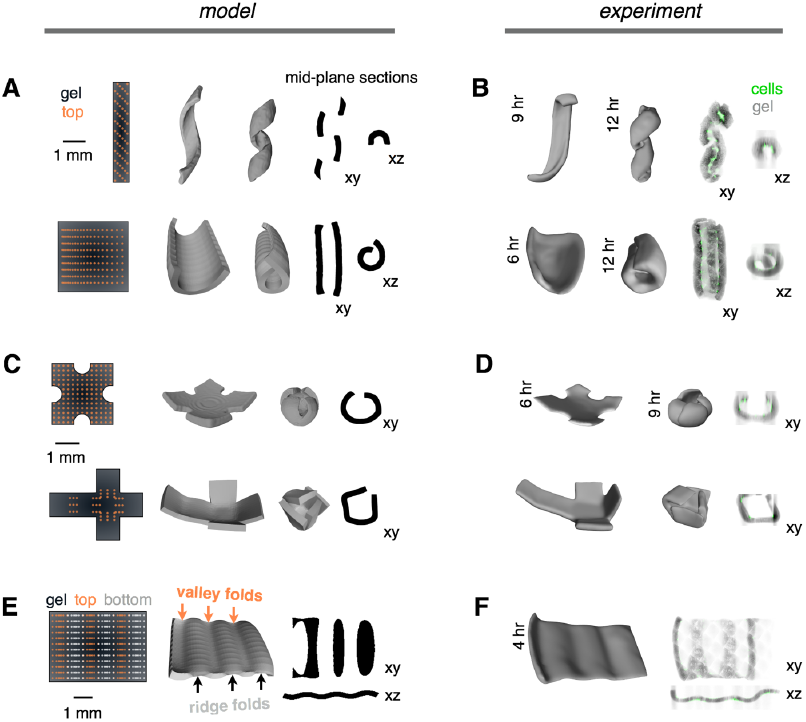
Networks of Mechanically Active Mesenchymal Condensates Program the Autonomous Folding of Diverse 3D Tissue Architectures. (A)(Left) DPAC blueprints of MEF cluster positions in x and y for a coiled object (top) and rolled tube object (bottom). (Center) Snapshots from the FEM showing intermediates along the predicted folding trajectories. (Right) Mid-plane sections from the FEM snapshots. (B)Shell surfaces and mid-plane sections of reconstituted tissues corresponding to the model objects in (A) at two imaging timepoints. Shell surfaces were reconstructed from confocal micrographs (MEFs, green; AF555-collagen I, gray). (C),(D) Spherical and cubic FEM objects and corresponding reconstituted tissues, imaged and analyzed as in (A) and (B). (E)(Left) DPAC blueprint encoding an opposing curvature motif incorporates MEF clusters on both the top (orange) and bottom (white) surfaces of the gel. (Center) FEM showing predicted opposing ridge and valley folds, and (right) mid-plane sections. (F)Shell surface and mid-plane sections of a reconstituted tissue corresponding to the model in (E).

Tubular structures are ubiquitous in the body, where ducts and vessels serve to carry fluids within and among organ systems. We reasoned that we could roll a tube from an initially planar sheet by combining anisotropic and compound curvature motifs. Specifically, a gradient in curvature along the horizontal axis was encoded using MEF clusters patterned horizontally with log-distributed spacings from 80 to 250 microns, each row being set apart by 300 microns along the vertical axis (movie S4). The folded tube was hollow along its entire length, with an inner diameter of 550 microns +/- 9% CV lengthwise (555 microns +/- 4% by FEM).

A number of tissues incorporate neighboring folds with opposing polarity. For example, the folds of the cerebral cortex and developing chick gut pass through intermediates having a locally zig-zagged architecture in which three folds of positive curvature and another of negative curvature converge at a point. We reconstructed this pattern by incorporating an opposing curvature motif into a fourfold-vertex shape (movie S5). We found that these more complex opposing fold patterns were surprisingly robust to errors in cell patterning. More specifically, folding was robust to “pop-through” defects so long as the cluster densities encoding adjacent folds were comparable (Method Details, Figure S6A-C).

The folding of flat surfaces into 3D shapes has fundamental physical constraints, since “non-developable” surfaces that are curved in more than one direction at a given location require that the material stretch or compress (Modes et al., 2010). Residual stresses not relieved by deformations in the material can lead to unintended folding outcomes (Armon et al., 2011; Kim et al., 2012). In vivo, these residual stresses could be relieved by additional out of plane buckling, or by in-plane changes in cell shape, size, intercalation geometry, proliferation, or ECM compaction (Humphrey and Dufresne, 2014; Legoff et al., 2013). Indeed, when we prepared a non-developable fold consisting of two adjacent isotropic folding motifs of opposite polarity in a single ECM gel, local folds emerged as expected, but additional buckling events occurred, consistent with the FEM-predicted accumulation of unresolved in-plane stresses (Figure S5D,E). In such cases, targeted modification of tissue boundary conditions by strategic cutting of surfaces prior to folding can enable the construction of more complex shapes without additional non-programmed buckling events, for example spherical and cubic tissues (as in the Kirigami art form) (Sussman et al., 2015; Zhang et al., 2015). We built both tissue architectures using laser microdissection of gel substrates prior to folding (Figure 5C,D, Figure S5F,G); in the latter case relying on anisotropic curvature at cube creases to actuate folding.

Finally, we combined anisotropic, compound, and opposing curvature motifs to generate a corrugation of pitch 1.6 mm and amplitude 130 microns (1.6 mm and 115 microns by FEM, Figure 5E,F, Figure S5H,I). These corrugated objects model periodic curvature patterns seen in vivo, for example at the dermal-epidermal junction in the skin, in the trachea, and a transient folding pattern of the luminal surface of the chick gut (Shyer et al., 2013).

### Reconstituting Tissues with Multiple Cell Types and Tessellated Curvature Motifs

The process of morphogenesis often results in the tessellation of architectural motifs, such as the repeated copies of nephrons in the kidney and the alveoli of the lung. We reasoned that the simple folding patterns described above could be tiled as repeating subunits to construct reconstituted tissue architectures of even greater size and complexity. Inspired by the zig-zag-shaped luminal surface of the embryonic day 13-16 (E13-16) chick gut (Shyer et al., 2013), we designed a tessellation of the four-fold junction in analogy to the Miura origami fold (Figure 6A). The Miura fold has several unusual geometric features, including the capability to be fit generically to complex target surfaces (Dudte et al., 2016). Finite element modeling of the Miura design predicted a folding trajectory with similarity to the folding pattern of the chick gut, but directed by cell-generated forces of an entirely synthetic design (Figure 6B; movie S4B). Consistent with the model, the reconstituted Miura tissue autonomously emerged from a 6 × 8 mm flat sheet to a 4 × 6 mm zig-zag structure at 15 hr with all 31 folds having the correct orientation, and similar periodicity and amplitude as those in the FEM.

**Figure 6.**
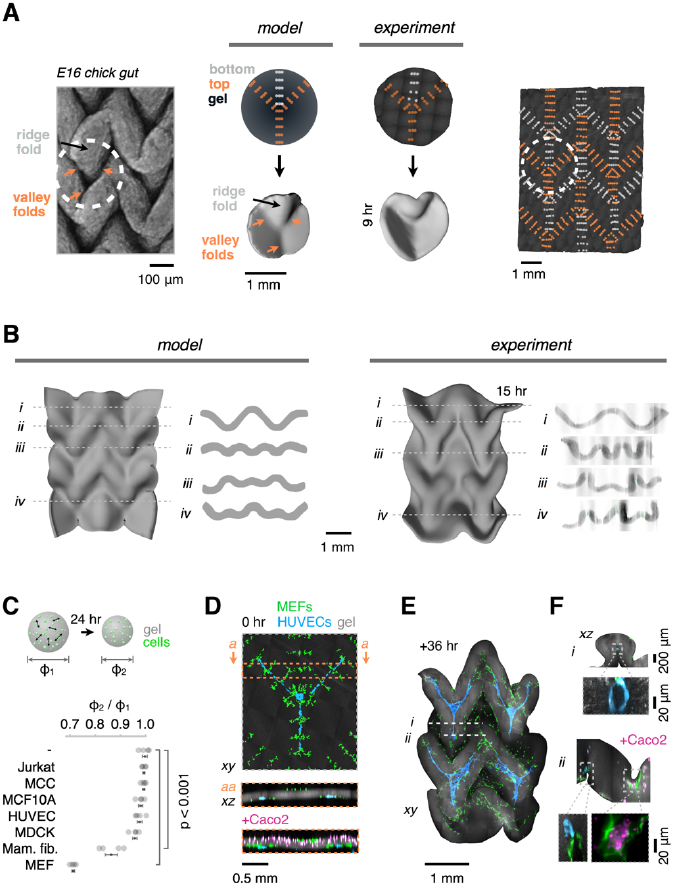
Mesenchymal Condensates Drive the Autonomous Folding of Tessellated Tissue Patterns Incorporating Multiple Cell Types. (A)(Left) Macro-confocal micrograph of Ethidium Bromide-stained embryonic day 16 (E16) chick gut lumen exhibiting a tessellated four-fold vertex pattern that incorporates three valley folds and one ridge fold converging on a single point. (Middle) FEM and as-printed DPAC blueprints, FEM snapshot, and shell surface of a reconstituted four-fold vertex tissue (AF555-labeled collagen I in gel, dark gray). (Right) As-printed DPAC blueprint encoding a tesselated architecture similar to the chick gut lumen and Miura origami fold. (B)(Left) FEM snapshot and cross-sections of the Miura object. (Right) Shell surface and cross-sections of the corresponding reconstituted tissue after 15 hr in vitro. (C)The shrinkage in diameter of collagen I-containing gels droplets (ɸ_2_/ɸ_1_) is quantified as an assay for the relative traction capacity of different cell types (mean ± SEM, *n* = 4, one-way ANOVA with Holm-Sidak’s multiple comparisons test). (D)Confocal micrographs and sections showing the DPAC output for a Miura folding pattern as in (B), but incorporating human umbilical vein endothelial cells (HUVECs) patterned as 3-pronged cords at programmed folds and/or Caco2 cells distributed uniformly at the top surface of the gel. (E)Maximum intensity confocal projection of the folded architecture of HUVEC-containing gel after 36 hr in culture. (F)(Top, *i*) Representative confocal cross-sections of the object in (E) showing lumenized HUVEC cords enveloped by Miura folds and (bottom, *ii*) Caco2 cells in a different HUVEC-containing Miura tissue deposited as in (D, bottom). Caco2 cell clusters form atop contractile fibroblasts within concave folds.

A remarkable aspect of tissue folding processes during development is that curvature trajectories are generally robust, even within microenvironments with complex cellular compositions (Nelson, 2016; Savin et al., 2011). Such robustness would also be important for engineering tissue folding, particularly when incorporating the additional cell types necessary to build a functional tissue. We therefore tested the robustness of mesenchymal condensation-driven folding in reconstituted tissues incorporating other cell types as “passengers”. We reasoned that a given folding trajectory predicted by FEM would not be disrupted if the rate of strain that passenger cells induced on their surrounding ECM was significantly lower than that of condensing mesenchymal cells. We therefore screened 7 cell types for their ability to contract ECM droplets, finding that MEFs and primary human mammary fibroblasts contracted the ECM at a much higher rate than other common cell types, including endothelial and epithelial lines (Figure 6C). These data suggested that the latter cell types, themselves critical components of most tissues, would not interfere with folding trajectories dominated by the properties of the mesenchyme. Additionally, our ability to include additional cell types in juxtaposition with condensing mesenchyme raised the intriguing possibility that the behavior of passenger cells would themselves be altered by the dynamics of the surrounding tissue architecture as it folded over the course of the experiment (Mammoto et al., 2013; Shyer et al., 2015).

To test these hypotheses we patterned multiple cell types in reconstituted tissues directed to fold into the Miura pattern previously described (Figure 6D-F, movie S6). 3-pronged HUVEC cords were positioned underneath incipient folds; while Caco2 cells (a colon carcinoma cell line) were deposited on top by embedding them uniformly near the tissue surface. We found that the folded shapes of these passenger cell-laden Miura tissues were similar to those predicted by FEM, confirming that the properties of the condensing MEFs dominated folding trajectories (Figure 6B,E). We additionally found that HUVEC migration was biased along incipient folds, suggesting that emerging tissue topography and ECM compaction induced by tissue folding feed back on the behavior of passenger cells (Method Details, Figure S6D,E). At later timepoints, the HUVEC cords became fully encapsulated within the zig-zag folds, and were lumenized across 100-200 micron tracts after 36 hr. Finally, we found that Caco2 cells, which form 3D cysts in matrigel (Ivanov et al., 2008), became concentrated at the base of valley folds on top of the network of fibroblasts. This completely synthetic folded architecture had a gross similarity to the developing small intestine (Walton et al., 2016a), indicating that mesenchymal compaction of the ECM may represent a general strategy for engineering folds into tissues with complex cellular compositions.

## DISCUSSION

Animal development involves the step-wise elaboration of tissue structure at multiple length-scales. Each step of morphogenesis acts upon and remodels the architecture formed in preceding steps. Thus, tissues are inherently imprinted with a developmental history that contributes to the anisotropy in their ECM, cell shape, and topography – architectural features that are critical for sculpting local cell-fate decisions, and for directing subsequent self-organization processes that determine tissue function (Brownfield et al., 2013; Cerchiari et al., 2015; Engler et al., 2006). Mesenchymal condensation is an example of a core vertebrate developmental program underlying this imprinting process, acting at keys steps of development in multiple tissues to remodel the topography, ECM anisotropy, and ECM density at the interface of the epithelium and mesenchyme (Kim et al., 2015; Li et al., 2015; Mammoto et al., 2013; Oster et al., 1983; Walton et al., 2012). The events that coincide with the emergence of curvature during mesenchymal condensation are complex, involving changes in both the mechanics and paracrine signaling microenvironment between multiple cellular components in each layer (Eames and Schneider, 2005; Varner and Nelson, 2014a; Walton et al., 2012). However, the contribution of the mechanics of the mesenchyme in these processes has not been studied in detail, and the onset of curvature is hypothesized to be driven primarily by forces generated by epithelial cell behaviors such as migration, localized growth, or shape changes (Lecuit et al., 2011; Panousopoulou and Green, 2016). We find that in the context of a loose and fibrous ECM composite, the mechanics and dynamics of condensing mesenchymal cells are sufficient to explain a variety of shape transitions in nearby tissue interfaces. In these cases, the mesenchyme behaves like an active composite material, with cells straining and compacting the ECM, aligning ECM fibers between regions of compaction, and encoding forces in the material along regions of maximum fiber alignment. These forces lead to bending of the material at tissue interfaces along trajectories that can be predicted using finite element modeling.

Importantly, the predictable relationship between strain and curvature observed in these cell-ECM composites allowed us to program the autonomous folding of tissues into a variety of 3D architectures bearing striking similarity to structures found *in vivo*, as well as into entirely novel geometries. The folding process is analogous to the autonomous folding of abiotic materials into complex shapes (Holmes et al., 2011; Kim et al., 2012; Na et al., 2014; Sydney Gladman et al., 2016; Tallinen et al., 2016).Moreover, the self-organizing and dynamic properties of a mesenchymal cell-ECM composite bear striking similarity to phenomena observed in reconstituted actomyosin networks (Köster et al., 2016; Linsmeier et al., 2016), suggesting these active materials may adhere to common physical principles.

One key aspect of our model is that it does not invoke any physical property of overlying tissue layers, such as an epithelium. However, it predicts that these properties would affect the magnitude and polarity of tissue folding. Here, reconstituted tissues and finite element models treat the overlying material as having negligible bending modulus. Thus, a condensation near the upper surface of reconstituted tissues always forms a region of concave curvature, and we leverage this idea to pattern complex folds by placing condensing cells at either the upper or lower surfaces of an ECM slab. However, if the overlying material has a higher bending modulus than the mesenchyme, modeling predicts an inversion of the curvature direction, converting an invagination into an evagination (Figure S1). The model further suggests a coincident lateral compaction of the overlying layer during a condensation event, forming a placode-like structure (Oster and Alberch, 1982). These studies leave open an intriguing possibility: that paracrine signaling originating in the mesenchyme could serve to set the mechanical properties of an overlying epithelium, thereby determining the direction and magnitude of folding during a condensation event in vivo. Such a view of mesenchymal-epithelial interaction could explain how different combinations of epithelium and mesenchyme transition to markedly different tissue architectures through an interplay between tissue mechanics and paracrine signaling.Combined with the established roles of the epithelium in tissue buckling, our results suggest that a combination of mechanically active tissue components could collaborate to initiate and reinforce the pattern, polarity, and magnitude of tissue folding (Hirashima, 2014; Lecuit et al., 2011; Nelson, 2016; Odell et al., 1981; Oster and Alberch, 1982; Savin et al., 2011; Shyer et al., 2013; Tallinen et al., 2016; Varner et al., 2015). These possibilities warrant further investigation.

Apart from its relevance to developmental biology, our study raises the possibility that dynamic control over both the material and physical properties of cell-ECM composites is readily achievable. In this view, building tissues de novo is a 4D process – where initial tissue structures and boundary conditions are assembled in 3D, but evolve in time across multiple length-scales according to specific developmental principles, converging ultimately on a new 3D structure with more defined and life-like structural features. This approach could significantly improve the structure, maturation, and vascularization of organoid tissue models at mesoscale (Lancaster and Knoblich, 2014; Takebe et al., 2015), and should be incorporated as a design criterion for 3D printed tissues. We believe these efforts have important implications for the engineering of *in vitro* models of disease, for regenerative medicine, and for future applications of living active materials such as in soft robotics.

## AUTHOR CONTRIBUTIONS

A.J.H. and Z.J.G. conceived the project. A.J.H. built reconstituted tissues. A.J.H. and M.T.L. imaged and characterized reconstituted tissue morphology. A.J.H. and J.Z. developed finite element models. M.C.C. did gel contraction assays. A.J.H. and H.M. did mouse gut assays. A.J.H., D.C., and Z.V. did chick skin and gut assays. All authors analyzed data and wrote the manuscript.

## ACKNOWLEDGMENTS

We thank J. Liu, A. Paulson, C. Krishnamurthy, J. Farlow, M. LaBarge, and J. Debnath for providing reagents and cells; M. Chung for technical assistance with chick gut dissection and imaging; A. Long and S. Dumont for discussion and assistance with laser ablation studies; A. Cerchiari, M. Thompson, and J. Spence, for critical discussion. This work was funded through a Jane Coffin Childs postdoctoral fellowship to A.J.H., NIH grants R01 DE016402 and S10 OD021664 to R.A.S., the Department of Defense Breast Cancer Research Program (W81XWH-10-1-1023 and W81XWH-13-1-0221 to Z.J.G.), the NIH Common Fund (DP2 HD080351-01 to Z.J.G.), the NSF (MCB-1330864 to Z.J.G.), the UCSF Program in Breakthrough Biomedical Research, and the UCSF Center for Cellular Construction (DBI-1548297), an NSF Science and Technology Center.

## METHOD DETAILS

### Cell Culture, Fluorescence Labeling, and Viability

Mouse embryonic fibroblasts (MEFs, gift of Jay Debnath, UCSF) expressing maxGFP via pSicoR-Ef1a-maxGFP-Puro (gift of Justin Farlow, Serotiny Bio), MCF10A cells expressing H2B-GFP (Liu et al., 2012), MCF10AT (Liu et al., 2012) (mesenchymal-like carcinoma cells, MCCs, bulk population and clones) expressing H2B-GFP and constitutively active H-Ras^V12^, human umbilical vein endothelial cells (HUVECs, Lonza) expressing mCherry after transduction with lentivirus made with pSicoR-Ef1a-mCh-Puro (Addgene 31845), human 998 mammary fibroblasts (gift of Mark LaBarge, LBNL) expressing maxGFP, Madin-Darby canine kidney epithelial cells (MDCK, UCSF cell culture facility), Jurkat immortalized T-cells (American Type Culture Collection), and Caco2 human epithelial colorectal adenocarcinoma cells (UCSF cell culture facility) were cultured on polystyrene plates and flasks (Corning). MCCs and MCF10A cells were cultured as previously described (Debnath et al., 2003). HUVECs were maintained in EGM-2 medium (Lonza). Jurkat cells were maintained in Roswell Park Memorial Institute (RPMI) medium with 10% fetal bovine serum (FBS). MEFs, 998 fibroblasts, MDCKs and Caco2 cells were maintained in Dulbecco’s modified Eagle’s medium (DMEM) with 10% FBS and 1x non-essential amino acids. Staurosporine (Sigma-Aldrich S5921) was prepared by serial dilution from a 10 mM DMSO stock. Cell lines without endogenous fluorescent markers were labeled with CellTracker Deep Red or Blue CMHC dyes according to manufacturer protocols (ThermoFisher Scientific C34565 and C2111 respectively). Cell viability was measured via microplate-based PrestoBlue assay according to manufacturer instructions (ThermoFisher Scientific A13261).

### Reconstituted Tissue Fabrication

Microfluidic flow cells were constructed by sandwiching aldehyde-silanized glass slides (Schott 1064874) against a polydimethylsiloxane (PDMS) membrane gasket (0.01” thick, SSP M823) cut with a craft cutter (Silhouette). Prior to sandwiching, through-holes in the top slide were made using 20 passes of a 50 W etching laser at 100% power, 15% speed, 350 pulses per inch (VLS3.5, Universal Laser Systems).Fiducial marks were etched into both glass slides to aid alignment by light microscopy before cell patterning. Etched slides were used as substrates for DNA-programmed assembly of cells (DPAC), as detailed previously (Todhunter et al., 2015). To summarize, after etching, amine-modified oligonucleotides were printed onto the slides using a microfluidic cantilever (NanoEnabler, Bioforce Nanosciences) and covalently attached by reductive amination. Printing locations were specified in bitmap files.Slides were treated with hydrophobic silane and blocked for 1 hr in 3% bovine serum albumin in PBS before being assembled against the PDMS gasket. Immediately prior to sandwiching, PBS was added to gaps in the gasket to prime flow cells and reduce the tendency to trap bubbles between the slides. Slight pressure was applied to the sandwich using a microarray hybridization cassette (AHC1X16, ArrayIt). The positions of through-holes in the top glass slide and voids in the PDMS gasket were matched to the dimensions of the cassette such that 16 flow cells could be independently addressed through pairs of through-holes.

Cells to be assembled on the top and bottom flow cell walls were lifted from plates using a PBS wash followed by 0.05% trypsin and labelled with lipid-modified oligonucleotides as previously described (Todhunter et al., 2015) (with the exception of Caco2 cells, see below). With the flow cell cassette on ice, cells in suspensions of ~ 10 × 10^6^ cells/ml were introduced to flow cells by gentle pipetting on top of one of the pair of through-holes and adhered to DNA spots on the glass. A further round of cellular assembly was used to generate clusters of 5-8 cells at each DNA spot. After cell patterning, liquid gel precursor was introduced in two aliquots of 20 μ l per flow cell and the cassette placed at 37 °C for 20 min to set the precursor. Reconstituted tissue gels consist of a composite of fluorescently-labeled collagen I fibers in matrigel. To prepare the gel precursor, 200 μ l of ~ 8.5 mg/ml rat tail collagen I in 0.02 M acetic acid (Corning 354249) was labeled using 5 μ l of 1 mg/ml Alexa Fluor 555- or 647-NHS ester in DMSO (ThermoFisher Scientific A20009 and A20006; chosen to avoid spectral overlap with cell labels in a given experiment) that was added immediately prior to neutralization with 10 μ l 20x PBS and 4 μ l 3 M NaOH on ice. After 10 min on ice, 70 μ l of this collagen stock was added to a second stock consisting of 415 μ l of ~ 9 mg/ml matrigel (Corning 354234) and 15 μ l of Turbo DNase (ThermoFisher Scientific AM2238). This 500 μ l precursor solution was sufficient to build reconstituted tissues in 8-10 flow cells.

The flow cell cassette was then disassembled gently, and the slide sandwich submerged in the appropriate cell media at room temperature. A razor blade was used to gently pry apart the glass slides. Reconstituted tissues consisting of cell clusters carried along with the ECM gel typically floated spontaneously into the media or could be gently detached from one of the glass slides with a micro-spatula (Fine Science Tools 110089-11). Floating tissues were then manually cut out using either a biopsy punch or razor blade, or by laser microdissection (Zeiss PALM MicroBeam). Finished tissues were transferred to glass coverslip-bottomed 24-well plates (Greiner) using a P1000 pipet trimmed to a ~ 7 mm diameter. If reconstituted tissues were intended to undergo folding, the glass in each well was coated with 1% agarose in PBS prior to adding tissues to prevent them from adhering. For imaging studies of collagen strain/alignment, cell migration, and non-folding controls, or prior to microdissection, reconstituted tissues were encouraged to adhere to the bottom of coverslip wells by 10 min 37°C incubation in a semi-dry state (with media temporarily withdrawn).

Rather than being assembled by DPAC, a semi-confluent layer of Caco2 cells was assembled at the lower tissue surface by pre-mixing them at 4 × 10^6^ cells/ml in gel precursor such that they settled onto the bottom of the flow cell prior to setting at 37°C.

Reconstituted tissues that contained HUVECs were cultured in EGM-2 with 200 ng/ml each of IL-3, stromal-cell derived factor 1a (SDF-1a) and stem-cell factor (SCF) to encourage lumenization. Tissues with a single passenger cell type were cultured for 12 hours in the appropriate fibroblast medium, and transferred to the passenger cell type’s medium thereafter. 3-cell type reconstituted tissues (containing MEFs, HUVECs, and Caco2 cells) were similarly transferred to 50:50 HUVEC:Caco2 media after 12 hours. We anticipate that significant optimization of media conditions will be required depending on the targeted endpoints relevant to cell types of interest.

### Droplet Contraction

Rapid screening for cell contractility was performed by setting 1 μ l droplets of ECM gel containing 10^6^ cells/ml on coverslip-bottomed microwells at 37°C for 20 min. Media was then added and droplets detached by gentle pipetting prior to timelapse imaging.

### Measuring ECM Gel Strain by Single Cells

ECM strain in the vicinity of single cells was determined by particle image velocimetry (PIV ImageJ plugin (Tseng et al., 2012)). 30 μ l aliquots of reconstituted tissue gel precursor with 1 µm red fluorescent polystyrene beads at 8 × 10^6^ per ml (ThermoFisher Scientific F13083) and unlabeled collagen fibers were set in wells of a 96-well plate at 37°C for 20 min. 3,000 cells per well were then settled on the gel underlay and imaged every 60 min by confocal microscopy for bead displacement.

### Collagen Fiber and Strap Orientation Analysis

Collagen fiber orientation and FEM edge orientation were measured using OrientationJ in ImageJ (Pü spö ki et al., 2016). The absolute value of pixels in orientation images was taken to put orientations in the 0-90° range. Orientation images were then smoothed using Gaussian blur on a length-scale of ~ 0.1 times the cell cluster spacing to enable interpretation. Average projections of aligned orientation images were used to visually summarize replicates.

Collagen strap orientation was determined in ImageJ by manual annotation. Straps that showed collagen fluorescence above a signal-to-noise ratio of 3 (measured from fluorescence profiles taken orthogonal to strap axes) were included in orientation plots.

### Laser Ablation of ECM Gel

Gel rebound was measured 30 min after ablation of ~ 100 µm x 10 µm tracks by laser microdissection (Zeiss PALM MicroBeam, Method Details). Laser power was adjusted in control regions distant from cell clusters to ensure cutting through the entire gel thickness.

### Reconstituted Tissue Imaging

3D timelapses of reconstituted tissue folding were recorded at 37 °C and 5% CO_2_ with a 10x objective via tiled confocal microscopy. Images were acquired through Zeiss Zen 2012 software with 30 µm z slice spacing using a Zeiss Observer.Z1 with a Yokogawa CSUX1 spinning disk and Photometrics Evolve 512 EMCCD camera.

### Spatial Reconstruction of Reconstituted Tissues from Confocal Data

Confocal image stacks were segmented in ImageJ either manually or by semi-automated thresholding. Binary image stacks were then read into MATLAB (MathWorks) and converted to isosurface meshes via marching cubes. Meshes were smoothed and face colors assigned to local mean curvatures using custom scripts.

Curvatures of isotropic reconstituted tissues were generated by radially reslicing and thresholding image stacks in ImageJ after manually picking the center of tissues to serve as origin points. Radial slices were read into MATLAB and least-squares fitted to circles using custom scripts. Anisotropic curvatures were similarly generated for the indicated cutting planes and folds.

### Example Reconstituted Tissue Immunofluorescence Protocol

Reconstituted tissues were transferred to a fresh 24-well plate and fixed in 2% paraformaldehyde for 45 min at room temperature. All pipetting was done while observing by light microscopy to avoid mechanical disruption with the pipet tip.Reconstituted tissues were washed three times with 100 mM glycine in PBS for 20 min per wash and permeabilized in 0.5% Triton X-100 in PBS for 15 min, all at room temperature; and blocked overnight at 4oC in 0.1% bovine serum albumin, 0.2% Triton X-100, 0.04% Tween-20, 10% goat serum (ThermoFisher Scientific 16210064) including a 1:50 dilution of goat anti-mouse IgG fab fragment (Jackson ImmunoResearch 115-007-003) in PBS. Reconstituted tissues were probed with primary antibodies in blocking buffer overnight at 4°C and washed three times for 1 hour per wash in blocking buffer. This process was repeated for secondary antibodies. Reconstituted tissue were then imaged in FocusClear (CedarLane FC-101).

### Mouse Gut Explant Culture and Immunofluorescence

PDGFRα^EGFP/+^ mice were used for gut explant studies (Hamilton et al., 2003). Embryonic intestines were harvested in cold DPBS or BGJb medium. Embryonic day 13.5 intestines were cut into approximately 5mm segments and either fixed immediately or cultured on 8.0 μ m pore-size transwell membranes (Corning, 353182) following the protocol described by Walton and Kolterud (Walton and Kolterud, 2014). Explants were cultured at 37°C and 5% CO_2_ in medium with or without 30 μ M (-)-Blebbistatin (Sigma-Aldrich, B0560), with media changed twice daily.

Explants were fixed overnight at 4C in 2% paraformaldehyde in PBS. Whole mount immunostaining was performed as previously described by Hatch and Mukouyama (Hatch and Mukouyama, 2015). In brief, tissue were blocked with blocking buffer (10% heat-inactivated goat serum and 0.2% Triton X-100 in PBS) overnight at 4 °C. The tissues were then incubated overnight at 4°C with primary antibodies diluted in the blocking buffer and washed five times for 25 minutes per wash in washing buffer (2% heat-inactivated goat serum and 0.2% Triton X-100 in PBS). The same process was repeated for secondary antibodies. Tissues were then placed on glass slides and incubated with FocusClear for 10-40 mins at room temperature prior to confocal imaging (as for reconstituted tissues). Primary antibodies were rat anti-mouse E-cadherin (1:100, Abcam ab11512) and rabbit anti-mouse Collagen I (1:100, Abcam ab34710). Secondary antibodies were AlexaFluor 647-labeled goat anti-rat IgG and AlexaFluor 568-labeled goat anti-rabbit IgG (1:250, ThermoFisher Scientific A21247 and A11011 respectively).

### Chick Gut Explants

Fertilized white Leghorn chicken eggs were incubated, windowed, and otherwise handled as previously described (Schneider, 1999). Embryos were decapitated and dissected at the indicated embryonic day to remove the gut tube. Tubes were cut into rough 2 mm segments and then longitudinally to expose the luminal surface. Tissues were fixed in 4% paraformaldehyde, washed 3 times in PBS for 5 min per wash, stained in 2 μg/ml ethidium bromide in PBS for 15 min, and washed similarly, all at room temperature (Eames and Schneider, 2005). The luminal surface was imaged using a macro confocal microscope (Nikon AZ-C2Si) at 2x magnification.

### Chick Skin Explants

Chick embryos were similarly dissected by removing the prospective pteryla with fine forceps and pinning the skin to an underlay of 1% agarose. Explants were incubated in DMEM with 10% FBS and 1x pen/strep (with or without blebbistatin) at 37 °C and 5% CO_2_. Explants were fixed and prepared for immunofluorescence studies as for mouse gut explants.

### Finite Element Modeling

Reconstituted tissues were modeled using Kangaroo2, a position-based dynamics solver within the Rhinoceros Grasshopper (Robert McNeel & Associates) algorithmic modeling environment (Bender et al., 2014). We took a form-finding FEM approach to enable interactive, rapid prototyping. Cuboidal unit cells were constructed from quad mesh faces with two diagonals per quad. Unit cells were assembled to model 3D tissues at a scale of 100 µm per unit edge length and 2D tissues at 10 µm per unit edge length, balancing spatial resolution and simulation time. All edges were modeled as linear elastic elements with stiffness k1 and rest lengths equal to their initial lengths.Since cell cluster positions were specified at 10x higher resolution for DPAC than for mesh vertices in 3D models, *contractile nodes* to be specified on the upper or lower model faces were generated by rescaling and rounding DPAC cell cluster coordinates. The edges in the *xy* plane coincident with these nodes were specified as *active edges* with a > 3x-higher spring constant k_2_ relative to other edges in the spring network, and with rest length *s* manipulated by the user.

Models constructed in this way had two parameters – the stiffness of non-active edges k_1_, and the active edge rest length *s*. k_1_ = 100 Nm^-1^ was fitted such that 3D model isotropic caps with p_xy_ = 170 would reach the maximal observed curvature of ~ 3 mm^-1^ at *s* = 0 (Figure S4B); and k_1_ = 20 Nm^-1^ was fitted for 2D models by manually iterating it for *s* = 0 to approximate the local matrix orientation generated by single MEF clusters at a sampling distance *r* = 50 µm away and *t* ~ 6 hours (Figure S3B). These values of k_1_ were then used for all subsequent simulations. 3D tissues were typically simulated for 10 values of *s* in the range of 0-1 to produce a family of objects at intermediate folding states for each design. Simulated tissue models were exported from Rhino as stereolithography files, remeshed in Zbrush (Pixologic) to remove mesh face orientation defects, read into MATLAB, voxelized, converted to isosurface meshes, smoothed and face colors mapped to local curvatures using custom scripts.

Since *s* is not directly analogous to tissue folding time *in vitro*, models intended for visual and local curvature comparison to reconstituted tissues at a given time-point were chosen from object families based on qualitative similarity.

### Statistical Analysis

Data were analyzed for statistically significant differences between sample means via ordinary one-way ANOVA with Holm-Sidak’s multiple comparisons test in Prism 7 (GraphPad).

### Inferring Tension from Local Retraction after Ablation in Reconstituted Tissues

Since real-time imaging of retraction at the gel surface was limited by a mismatch in the appropriate length-scales of cutting and confocal microscopy, we made dissections and imaged resulting incisions within approximately 30 min. We expected that strain due to elastic tissue rebound would be fully realized within seconds (Bonnet et al., 2012), while continued strain due to active pulling at cell clusters would be limited to less than ~ 10 µm along collagen straps within 30 min (Figure S4E, movie S2A). We consider ablation of tissues to be analogous to severing an elastic spring under tension, discarding time-dependent dynamics from the Kelvin-Voigt model typically used to study viscoelastic retraction in live biological contexts (Bonnet et al., 2012; Kumar et al., 2006; Tanner et al., 2010). Therefore, the pre-tension force orthogonal to an ablated incision F∝ (L-L_o_), where L is the relaxed incision width at the gel surface after ablation, and L_o_ is the width of gel immediately destroyed by laser irradiation (in practice, the incision width in control gels lacking cell clusters).

### Scaling Analysis of Isotropic Reconstituted Tissue Curvature

We approximate reconstituted tissues as synclastic elastic plates, such that for small curvatures (deflections smaller than plate thickness), the strain in the *yz* plane at an arbitrary vertical distance *z* from the midplane of the plate is given by:

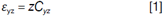

where *C*_*yz*_ is the curvature of the midplane parallel to the yz plane (Timoshenko and Woinowsky-Krieger, 1959). We assume that arrays of isotropic actuators on reconstituted tissues generate spatially homogeneous strains tangential to gel surfaces at a given timepoint *t* in proportion to their total number *N*_*c*_ such that:

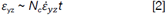

where 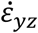 is the collagen matrix strain rate of a cell cluster. Experimentally measured values of 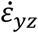 appear to be approximately constant over time for large ranges in local and global collagen gel deformation (Meshel et al., 2005). Thus, combining [1] and [2] suggests a first-order model of the form *C*_*yz*_ ∝ *N*_*c*_ to fit calibration data describing folding of isotropic reconstituted tissues at intermediate time-points.

### Fidelity of Anisotropic Folds

We hypothesized that cell-type specific differences in migration rate would predict the fidelity with which initial cluster positions encode anisotropic curvatures. We found that mesenchymal-like carcinoma cells (MCCs) migrated to a characteristic distance of half the cluster pair spacing *d*/2 of 80 µm in a characteristic time τ_d/2_ of 7.5 hr, while mouse embryonic fibroblasts (MEFs) reached this threshold in 11 hr (Figure S4I). Because the folding times for isotropic reconstituted tissues with grids of similar spacing were approximately 10 hr and 5 hr for MCC and MEF cell types, respectively, these data suggest that MCC clusters disperse from their initial positions before significant folding occurs, whereas MEFs do not. This dispersal of cells away from their intended positions during folding would tend to produce spatially uncontrolled strains, and thus uncontrolled curvatures.

To further investigate this hypothesis, we measured the angle at which tensile collagen straps formed between clusters (relative to the horizontal axis) in isotropic or highly anisotropic grids (Figure S4J). At 9 hours in culture, straps between clusters were primarily oriented towards nearest neighbors along 0 and 90 degree angles in isotropic grids, regardless of cell type. At 15 hours, however, strap orientation had become randomly distributed due to the disorganizing effect of cell migration away from initial cluster positions.

For anisotropic grids, straps between clusters were also primarily oriented towards nearest neighbors at 9 hr, focused at 0°along the horizontal axis. At 15 hours, however, straps were more frequently oriented away from this axis (i.e. were more disorganized) in anisotropic MCC grids when compared to MEF grids, consistent with extensive MCC migration away from initial cluster positions. These data suggest that a threshold in the ratio of cell strain rate to migration rate determines the extent to which a given cell type actuates reconstituted tissues with sufficient fidelity.

### Robustness of Adjacent Opposing Folds to Pop-Through

Of central concern for the design of adjacent folds of alternating negative and positive curvature (i.e. opposing folds) is their robustness to “pop-through” defects, in which out of plane deformations can emerge in the unintended orientation. Opposing folds can be built by placing cell clusters on opposite sides of the ECM sheet at spatially distinct regions in the *xy* plane. To study the robustness of our reconstituted tissues to such defects, we built four-fold vertex tissues that have three folds of the same orientation and one in the opposite orientation that converge at a single point. We hypothesized that for opposing folds to form successfully, they would need to be actuated by similar numbers of cell clusters so that the orientation of a particular fold would not be determined by a more radically curved neighboring fold. Indeed, a critical threshold at which the opposing fold formed in the incorrect orientation emerged when the number of clusters in the opposing fold was less than around half the average number in the two adjacent folds (Figure S6B). However, for values of this ratio above ~ 0.75, the opposing fold always popped into the correct orientation, demonstrating that adjacent opposing folds form robustly if spatial strain profiles are properly managed.

### Modulation of Passenger Cell Behavior

HUVEC cells patterned as a patch at the intersection of 3 folds in fourfold vertex reconstituted tissues migrated preferentially along nascent folds in comparison to non-folding control tissues (Figure S6D,E), perhaps responding to local collagen concentration and/or alignment orchestrated by fibroblasts at folds. These data suggest that reconstituted tissues can incorporate cells that generate the matrix strains and alignments necessary for folding, as well as other cell types that respond dynamically to these microenvironmental cues as folding progresses.

